# Not all voxels are created equal: reducing estimation bias in regional NODDI metrics using tissue-weighted means

**DOI:** 10.1101/2021.06.29.450089

**Authors:** CS Parker, T Veale, M Bocchetta, CF Slattery, IB Malone, DL Thomas, JM Schott, DM Cash, H Zhang, for the Alzheimer’s Disease Neuroimaging Initiative

**Affiliations:** Centre for Medical Image Computing, Department of Computer Science, UCL, London, UK; The Dementia Research Centre, Department of Neurodegenerative Disease, UCL Queen Square Institute of Neurology, UCL, London, UK; UK Dementia Research Institute at UCL, UCL, London, UK; Department of Brain Repair and Rehabilitation, UCL Queen Square Institute of Neurology, UCL, London, UK; Wellcome Centre for Human Neuroimaging, UCL Queen Square Institute of Neurology, UCL, London, UK

**Keywords:** Diffusion MRI, Microstructure imaging, Region-of-interest, Arithmetic mean, Tissue-weighted mean

## Abstract

Neurite orientation dispersion and density imaging (NODDI) estimates microstructural properties of brain tissue relating to the organisation and processing capacity of neurites, which are essential elements for neuronal communication. Descriptive statistics of NODDI tissue metrics are commonly analysed in regions-of-interest (ROI) to identify brain-phenotype associations. Here, the conventional method to calculate the ROI mean weights all voxels equally. However, this produces biased estimates in the presence of CSF partial volume. This study introduces the tissue-weighted mean, which calculates the mean NODDI metric across the tissue within an ROI, utilising the tissue fraction estimate from NODDI to reduce estimation bias. We demonstrate the proposed mean in a study of white matter abnormalities in young onset Alzheimer’s disease (YOAD). Results show the conventional mean induces significant bias that correlates with CSF partial volume, primarily affecting periventricular regions and more so in YOAD subjects than in healthy controls. Due to the differential extent of bias between healthy controls and YOAD subjects, the conventional mean under- or over-estimated the effect size for group differences in many ROIs. This demonstrates the importance of using the correct estimation procedure when inferring group differences in studies where the extent of CSF partial volume differs between groups. These findings are robust across different acquisition and processing conditions. Bias persists in ROIs at higher image resolution, as demonstrated using data obtained from the third phase of the Alzheimer’s disease neuroimaging initiative (ADNI); and when performing ROI analysis in template space. This suggests that conventional ROI means of NODDI metrics are biased estimates under most contemporary experimental conditions, the correction of which requires the proposed tissue-weighted mean. The tissue-weighted mean produces accurate estimates of ROI means and group differences when ROIs contain voxels with CSF partial volume. In addition to NODDI, the technique can be applied to other multi-compartment models that account for CSF partial volume, such as the free water elimination method. We expect the technique to help generate new insights into normal and abnormal variation in tissue microstructure of regions typically confounded by CSF partial volume, such as those in individuals with larger ventricles due to atrophy associated with neurodegenerative disease.

## 1. Introduction

Neurite orientation dispersion and density imaging (NODDI) (Zhang, H. et al 2012) is a widely used approach for estimating microstructural properties of tissue using diffusion weighted magnetic resonance imaging (DWI) (Alexander et al 2019). NODDI estimates the density and orientation dispersion of neurites, two key aspects of neurite morphology. These tissue metrics, termed neurite density index (NDI) and orientation dispersion index (ODI), relate to the density and structural organisation of axons in white matter and dendrites in grey matter that are essential for neural communication and provide useful biomarkers of brain function. Their changes have been linked to function in both healthy populations (Kunz et al 2014, Genc et al 2018, Mollink et al 2019) and in cases of diseases (Winston et al 2014, Broad et al 2019, Scahill et al 2020).

A common way of investigating NODDI tissue metrics is region-of-interest (ROI) analysis. This utilises descriptive statistics of NODDI metrics in ROIs to summarise microstructure in a region. These statistics may then be correlated to neurological phenotypes to make inferences about brain-phenotype associations. This approach has been applied in studies of normal development (Lynch et al 2020), aging (Kodiweera et al 2016) and in neurological diseases such as Huntington’s disease (Zhang, J. et al 2018, Scahill et al 2020), fronto-temporal dementia (Wen, J. et al 2019) and Alzheimer’s diseases (Colgan et al 2016, Slattery et al 2017, Parker et al 2018, Wen, Q. et al 2019).

Among the many approaches of imaging tissue microstructure with DWI, NODDI is one of the few that explicitly quantify the extent of CSF contamination, making it particularly suited for studying normal ageing and neurodegenerative diseases. NODDI employs a multi-compartment model to represent the signal from both CSF and tissue in a voxel, which enables quantification of NDI and ODI, microstructure parameters of the tissue space, that are free from CSF contamination. NODDI can therefore by used to investigate tissue microstructure abnormalities in conditions associated with brain atrophy without the confounding influence of CSF. This contrasts with other approaches, such as diffusion tensor imaging (Basser et al 1994), in which microstructure parameters in voxels contaminated by CSF are confounded (Metzler-Baddeley et al 2012).

However, this approach to removing CSF contamination presents a hitherto unrecognised problem in ROI analysis. Namely, the mean of NODDI tissue metrics in an ROI, such as the mean NDI or ODI, become biased when the ROI contains voxels with CSF partial volume. This is because conventional methods calculate the mean by averaging NODDI tissue metrics across all voxels in an ROI (Parker et al 2018, Zhang, J. et al 2018, Wen, J. et al 2019, Wen, Q. et al 2019, Andica et al 2020, Scahill et al 2020), which weight all voxels equally. In doing so, the variation in the amount of tissue present in a voxel due to CSF contamination is not accounted for. This can lead to a mis-estimation of the mean microstructure of the tissue across the ROI, which can be problematic for white matter tracts in periventricular regions whose border are particularly vulnerable to high CSF contamination. Given that ventricular enlargement is a prominent feature in individuals with Alzheimer’s disease (Schott et al., 2005), NODDI data from these patients are more likely to be affected by this issue.

To address this, we introduce the tissue-weighted mean, a new approach that aims to produce estimates of regional microstructure unbiased by the presence of CSF contamination. The key idea is to utilise tissue fraction metrics from NODDI to account for varying CSF contamination among ROI voxels. This is enabled by NODDI’s explicit representation of the tissue and CSF compartments. In contrast to the previous approaches that consider the influence of CSF partial volume on ROI means, such as the streamline density weighted average (Lynch et al 2020), this permits usage of the full tissue microstructural information across the ROI. Tissue-weighted means therefore account for CSF contamination while retaining the intent of the conventional means – which is to estimate the average microstructure metric across the brain tissue within the ROI.

In this work we provide a theoretical description and comparison between the conventional and tissue-weighted means using NODDI metrics. Our study aims to determine the prevalence across ROIs of estimation bias associated with the conventional mean, compare bias between healthy individuals and those with larger ventricles, and to assess the impact of applying the tissue-weighted mean to group studies. To do this, we apply the tissue-weighted mean in an exemplar study of white matter regional abnormalities in a cohort of healthy individuals and those with Young Onset Alzheimer’s Disease (YOAD), in which increased ventricular volume is a prominent feature of the disease (Drayer 1985). The conventional mean, tissue-weighted mean, bias and effect sizes are quantified in periventricular and non-periventricular ROIs. We also assess whether bias persists across alternative image resolutions and image analysis spaces.

The rest of this paper is organised as follows: we first formally define the tissue-weighted mean, before explaining the theory and methods used to quantify bias in conventional means. We then describe the methods for calculating conventional and tissue-weighted means of NODDI tissue metrics in a cohort consisting of cognitively healthy individuals and those with YOAD. Bias in conventional means and differences in effect sizes are compared and reported across white matter ROIs. Finally, we discuss our findings and their implications for studying brain regional microstructure.

## 2. Theory

### 2.1. NODDI model: voxel compartments and tissue parameters

NODDI adopts a multi-compartment model of the diffusion signal in each voxel (Zhang, H. et al 2012). The model assumes the DWI signal is a summation of signals from two primary compartments in a voxel: one representing free water and the other from tissue (Fig. 1). The free water fraction (FWF) parameter estimates the volume fraction of free water, contributed primarily from CSF. The tissue fraction (TF) parameter estimates the volume fraction of the tissue, TF=1-FWF.

**Figure 1.**
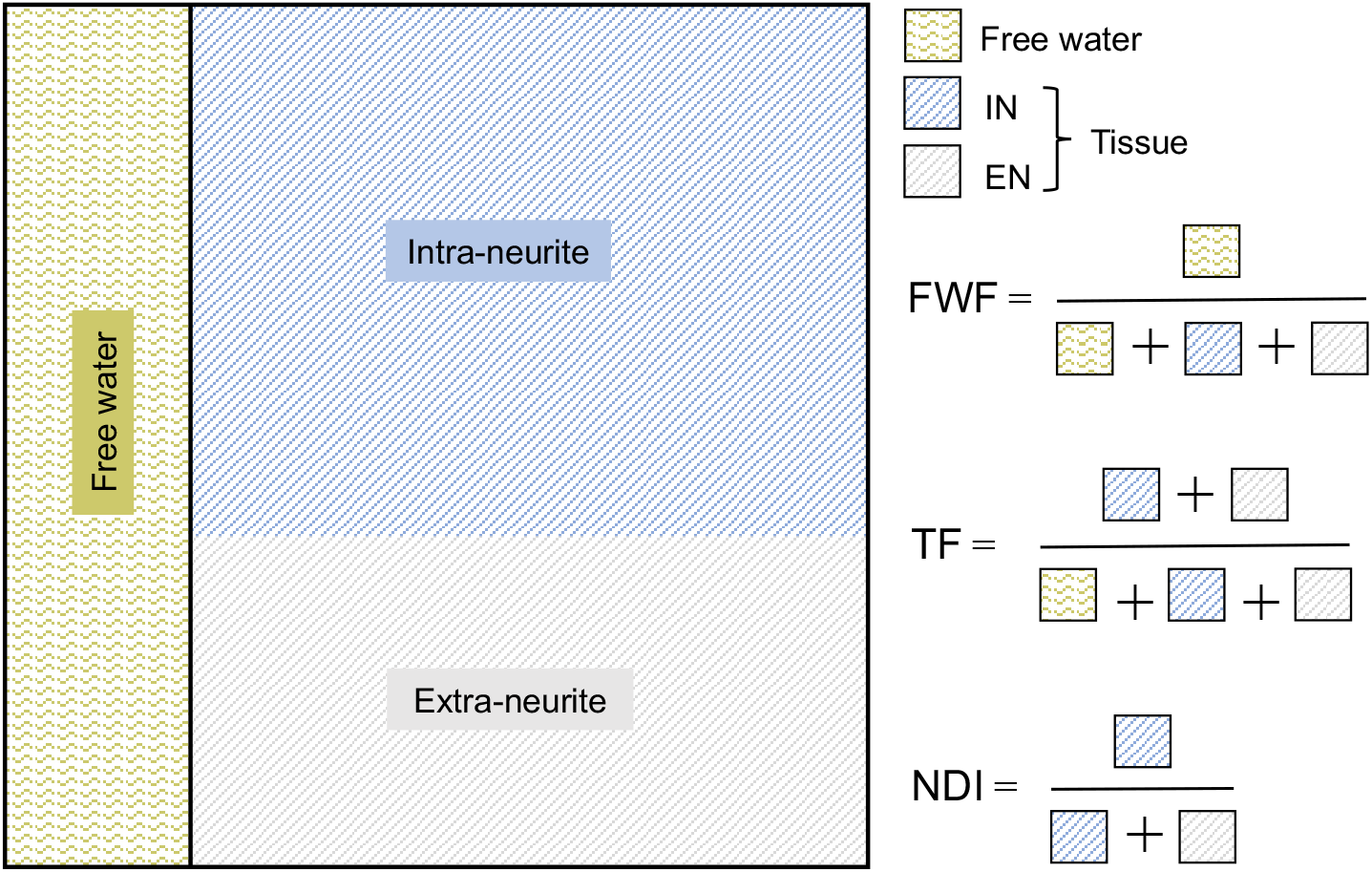
Graphical representation of the NODDI model. The model estimates the volume of the free water (area shaded with wiggly lines) and tissue (area shaded with straight lines) compartments within a voxel, parameterised as the free water fraction (FWF) and tissue fraction (TF), respectively. The tissue volume is further represented as two sub-compartments, each estimating the volume of the intra-neurite (IN) and extra-neurite space (EN) (lines coloured blue and grey, respectively). NDI estimates the density of intra-neurite space within the tissue and is equal to the relative volume fraction of the intra-neurite compartment. ODI is also a property of the tissue, representing the orientational distribution of the intra-neurite space.

Further parameters are derived corresponding to properties of the intra-neurite and extra-neurite space within the tissue component of the voxel. NDI provides a surrogate measure of the neurite density in the tissue compartment and ranges from 0 (low density) to 1 (high density). ODI estimates the dispersion of neurite orientations in the tissue and ranges from 0 (no dispersion) to 1 (fully dispersed).

### 2.2. Conventional mean

The conventional mean of NDI or ODI weighs each voxel equally and is calculated as the arithmetic mean. Let m denote the set of values for a NODDI tissue property of interest (i.e. NDI or ODI) in an ROI. Its arithmetic mean, denoted as 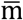, is then

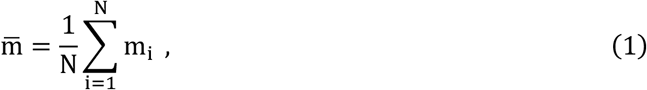

where i is the voxel index within the ROI, ranging from 1 to N, the number of voxels in the ROI. m_i_ is the NODDI tissue property for voxel i.

### 2.3. Tissue-weighted mean

The tissue-weighted mean instead calculates the mean of the metric across the tissue component of the ROI using a weighted average, with the weightings being the fraction of tissue in each voxel. The tissue-weighted mean of m, denoted as 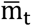, is thus:

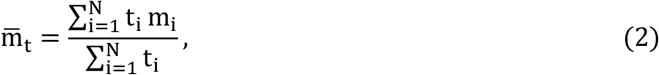

where t_i_ is the TF of voxel i within the ROI. This is equivalent to

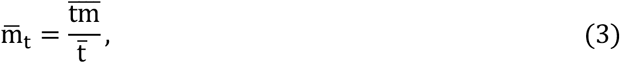

a more concise formulation derived by dividing both the numerator and denominator of equation (2) by N.

### 2.4. Bias in conventional means

Fig. 2 shows an illustrative example of calculating the two means in an ROI consisting of two voxels, each with different TFs. When calculating the conventional mean, NDI values in the voxel with lower TF are overweighted, resulting in a miscalculation of the tissue mean. In contrast, the tissue-weighted mean weights each NDI value by its voxel TF and gives correct calculations of the tissue mean. This demonstrates that the conventional mean is in general a biased estimate of the tissue-weighted mean.

**Figure 2.**
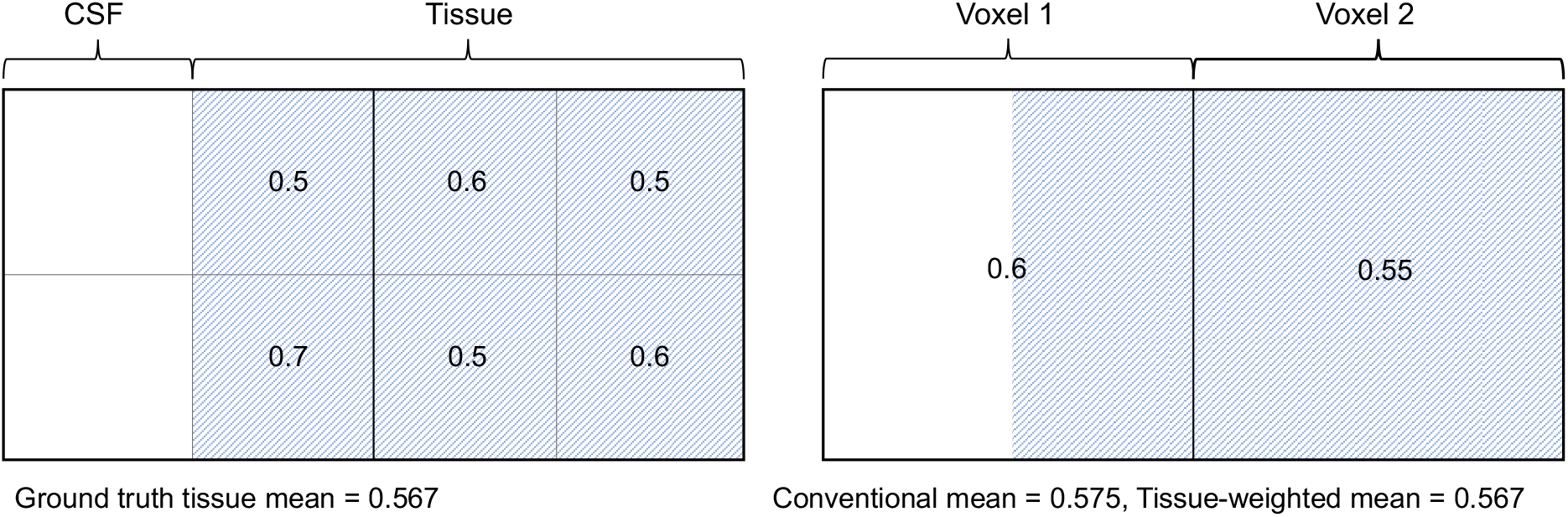
Illustrative case of estimation bias in the conventional mean in an ROI consisting of two voxels with different tissue volumes. Left shows an ROI covering an area of ground truth anatomy containing both CSF and tissue. Numbers in each sub-region show the local neurite density and are approximately normally distributed. Right shows NODDI tissue parameter NDI in two voxels covering the ROI. For each voxel, the NDI parameter describes the average neurite density of the tissue sub-regions that the voxel covers. In the presence of CSF partial volume, the conventional mean misestimates the ground truth mean by overweighting the NDI value in voxel 1, generating a bias of 0.008. Using estimates of the TF to weight the NDI in each voxel, the tissue-weighted mean correctly calculates the mean NDI of the tissue as 0.567.

To clearly show the relation between the two means, observe that the numerator in equation (3) can be written as

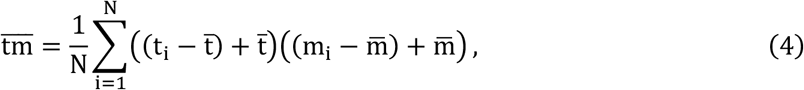

which simplifies to

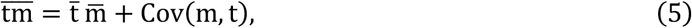

where Cov(m,t) is the covariance of m and t across the ROI. Here, Cov(m,t) is calculated without Bessel’s correction (with N instead of N-1 in the denominator), which is an unbiased estimate of the population covariance when N is large.

Dividing equation (5) by 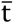 shows that the tissue-weighted mean is the conventional mean with an additional term, namely the covariance of m and t divided by 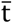. The difference between the means can be written as

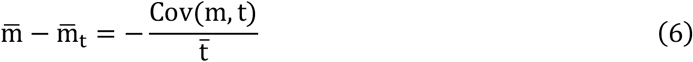

The term on the right-hand side is equal to the bias in the conventional mean. The two means are only equal when Cov(m, t) = 0. For a fixed covariance, the smaller the mean TF of the ROI, corresponding to higher partial volume, the larger the bias. The bias in the conventional mean is positive when there is a negative correlation between m and t and vice versa for positive correlation.

## 3. Materials and methods

### 3.1. Study participants

We analysed NODDI data from 21 control subjects and 30 patients that was acquired in a study of YOAD (Slattery et al 2017). Recruitment, diagnosis and exclusion criteria are described in the supplementary material (section S.2). Patient demographic characteristics are shown in Table S1. Ethical approval was obtained from the National Hospital for Neurology and Neurosurgery Research Ethics Committee and written informed consent obtained from all the participants.

### 3.2. Image acquisition

A multi-shell DWI sequence optimised for NODDI was acquired on a 3T Siemens Magnetom Trio scanner (Siemens, Erlangen, Germany) using a 32-channel phased array receiver head coil. DWI acquisitions consisted of a spin-echo echo planar imaging (EPI) sequence with EPI factor 96; TR=7000ms; TE=92ms; 55 interleaved slices with slice thickness 2.5mm; in plane FOV 240×240mm^2^ with resolution 2.5×2.5mm^2^; multi-slice acceleration factor 2; b-values=0 (n=13), 300 (n=8), 700 (n=32) and 2000 (n=64) s/mm^2^. Optimised gradient directions from the Camino software package generated using electrostatic energy minimisation were used (Cook et al 2007). Sequences utilized twice-refocused spin echo to minimize distortion effects from eddy-currents (Reese et al 2003). The total acquisition time was 16m13s.

T1-weighted images and B0 field maps were acquired to correct for susceptibility-induced off-resonance fields. T1-weighted images were acquired using a 3D sagittal MPRAGE volumetric sequence with TE=2.9ms; TI=900ms; TR=2200ms, matrix size 256×256×208 and isotropic 1.1×1.1×1.1mm^3^ voxels. For B0 field mapping, 2D dual echo gradient echo images were acquired using an EPI sequence with TEs=4.92/7.38ms and TR=688ms, matrix size 64×64×55 and resolution 3×3×3mm^3^.

### 3.3. Pre-processing

DWI non-brain voxels were removed by aligning the intracranial volume mask of the T1-weighted image to the DWI using SPM12 (Malone et al 2015). Inter-volume misalignment due to motion and image distortions due to residual eddy current-induced off-resonance fields were corrected using FSL eddy v6.0.2 (Jenkinson et al 2012, Andersson et al 2016). B0 maps were calculated from the unwrapped gradient echo phase images. Distortions in the DWIs due to susceptibility-induced off-resonance fields were corrected via a combined approach using the B0 maps and registration to the T1-weighted image (Daga et al 2014).

### 3.4. NODDI metrics

The NODDI model was fitted to the pre-processed DWIs using AMICO (Zhang, H. et al 2012, Daducci et al 2015), outputting parameter maps of NDI, ODI and FWF. TF maps were calculated from the FWF as 1-FWF using FSL fslmaths.

### 3.5. Atlas-based parcellation of white matter ROIs

NODDI maps in the native subject space were parcellated into forty-eight white matter ROIs defined in the John Hopkins University (JHU) white matter atlas (Mori et al 2008) using atlas-based parcellation. This was achieved by aligning the JHU atlas and subjects’ NODDI data via a bootstrapped population template. The alignment was also used to parcellate ROIs in the template space (see section 3.10).

The bootstrapped population template was created from the subjects’ DWI data (as in Zhang, J. et al 2018) using an iterative alignment algorithm (Guimond et al 2000, Zhang, H. et al 2007, Zhang, H. et al 2010) that employs linear and non-linear image registration in DTI-TK (Zhang, H. et al 2006). DTI-TK is a diffusion tensor (DT)-based registration, ranked the best of its kind (Wang et al 2011), which has been shown to improve alignment in white matter (Pecheva et al 2017) and reduce systematic errors compared to FA-based registration (Keihaninejad et al 2013). ROIs were propagated to the subject native space via the IIT atlas (Zhang, S. & Arfanakis 2011) and the bootstrapped population template. Propagation via the IIT atlas enables accounting for anatomical variation among JHU atlas participants.

Alignment between JHU and IIT atlases was implemented by registering their respective fractional anisotropy maps using FSL flirt and fnirt. IIT atlas and population template were aligned using linear and non-linear DT-based image registration in DTI-TK. Nearest neighbour interpolation was used to preserve the categorical nature of the labels.

ROIs were classified as periventricular (those sharing a border with ventricles, n=29) and non-periventricular (n=19) by manually inspecting ROIs overlayed on the between-subject average DT mean diffusivity maps in population template space. ROIs are shown in Fig. S1 and abbreviations are described in Table A1.

### 3.6. Conventional and tissue-weighted means

The conventional and tissue-weighted NDI and ODI means were calculated for each white matter ROI for each subject. Conventional means were computed using FSL fslstats. Tissue-weighted means were computed using the implementation available at https://github.com/tdveale/TissueWeightedMean, which is based on the alternative formula defined in equation (3). This formula can be readily implemented with FSL fslmaths and fslstats.

### 3.7. Mean tissue fractions

To understand the relation between the conventional mean and CSF partial volume contamination, mean TFs were analysed for all ROIs in control and YOAD subjects by calculating their between-subject mean and standard deviation. Differences in mean TF between groups were determined using two-tailed Welch’s t-tests, a variant of the students t-test that accounts for unequal variance between groups. A Bonferroni-corrected *p*-value threshold of <0.05 was applied to determine significant differences and control family-wise error (FWE) rate to <0.05. R v3.6.1 was used for these calculations and all subsequent analyses.

### 3.8. Bias in conventional mean

Bias in the conventional mean was estimated in all ROIs by subtracting the tissue-weighted mean from the conventional mean. The calculated bias was equal to its theoretically predicted value 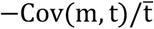 (see equation (6)).

Regional bias was summarised in control and YOAD subjects by their between-subject mean and standard deviation. Non-zero bias was determined using two-tailed one sample t-tests and the difference in bias between control and YOAD subject was determined using two-tailed Welch’s t-tests. Bonferroni-corrected *p*<0.05 were considered significant.

Pearson correlation was used to test for associations between mean TF and magnitudes of bias for control and YOAD groups and two-tailed Welch’s t-tests were used to test for different magnitudes of bias between periventricular and non-periventricular ROIs. In both cases *p*<0.05 was considered significant.

### 3.9. Bias at higher image resolution

We assessed whether bias exists in contemporary datasets with higher image resolution than that of the YOAD study. Multi-shell DWI, T1-weighted images and field maps for 75 healthy control subjects were obtained from the third phase of the Alzheimer’s Disease Neuroimaging Initiative (ADNI 3) (Mueller et al 2005, Weiner et al 2017, adni.loni.usc.edu). DWI and T1-weighted images were obtained from the earliest available visit for each subject and field maps closest to the DWI visit were selected. Subject characteristics are described in the supplementary material (section S.3).

The multi-shell DWI sequence, optimised for NODDI, was acquired on a 3T Siemens Prisma scanner (Siemens, Erlangen, Germany) using a 64-channel receiver head coil and consisted of a spin-echo EPI sequence with EPI factor 128; TR=3400ms; TE=71ms; 81 slices with slice thickness 2mm; in plane FOV 232×232mm^2^ with resolution 2×2mm^2^; multi-slice acceleration factor 3; b-values=0 (n=13), 500 (n=6), 1000 (n=48) and 2000 (n=60) s/mm^2^. Gradient directions were evenly spaced using an electrostatic repulsion algorithm (Caruyer et al 2013). The total acquisition time was 7m20s.

T1-weighted images and B0 field maps were obtained to correct for susceptibility-induced off-resonance fields. T1-weighted images were acquired using a 3D sagittal MPRAGE volumetric sequence with TE=2.95ms; TI=900ms; TR=2300ms, matrix size 240×256×176 and 1.05×1.05×1.2mm^3^ voxels. For B0 field mapping, 2D dual echo gradient echo images were acquired using an EPI sequence with TEs=4.92/7.38ms and TR=571ms, matrix size 78×78×54 and resolution 2.97×2.97×3.75mm^3^.

Imaging data underwent the identical pre-processing and ROI analysis steps as described in sections 3.2-3.8.

### 3.10. Bias in template space ROIs

As an alternative to native space ROI analysis (Oishi et al 2009, Zhang, S. & Arfanakis, K 2014, Wen, Q. et al 2019, Lynch et al 2020), template space ROI analysis is also common (Geng et al 2012, Kodiweera et al 2016, Zhang, J et al 2018, Schahill et al 2020). Hence we also examined bias in template space ROIs, by propagating JHU ROIs to the bootstrapped population template and repeating the analyses described in sections 3.6-3.8. The population template had voxel dimensions of (1.75 x 1.75 x 2.25) mm^3^.

### 3.11. Group differences comparison

To determine the implications of applying the tissue-weighted mean for group differences, Cohen’s d_s_ effect sizes (Cohen 1988), which describe the standardised mean difference between two groups of sample observations, were computed for each ROI (mean difference contrast of control minus YOAD). Effect sizes were compared between the conventional and tissue-weighted mean computed in the subjects’ native space. Two-tailed Welch’s t-tests were used to determine significant group differences. Bonferroni-corrected *p*<0.05 were considered significant.

## 4. Results

### 4.1. Mean tissue fractions

Mean TF varies between regions (Fig. 3) and as expected, tends to be lower in periventricular regions (Table A1), suggesting those ROIs experience higher CSF partial volume contamination. YOAD subjects’ ROIs tend to have lower mean TFs than control subjects, consistent with expected increases in CSF partial volume contamination due to atrophy.

**Figure 3.**
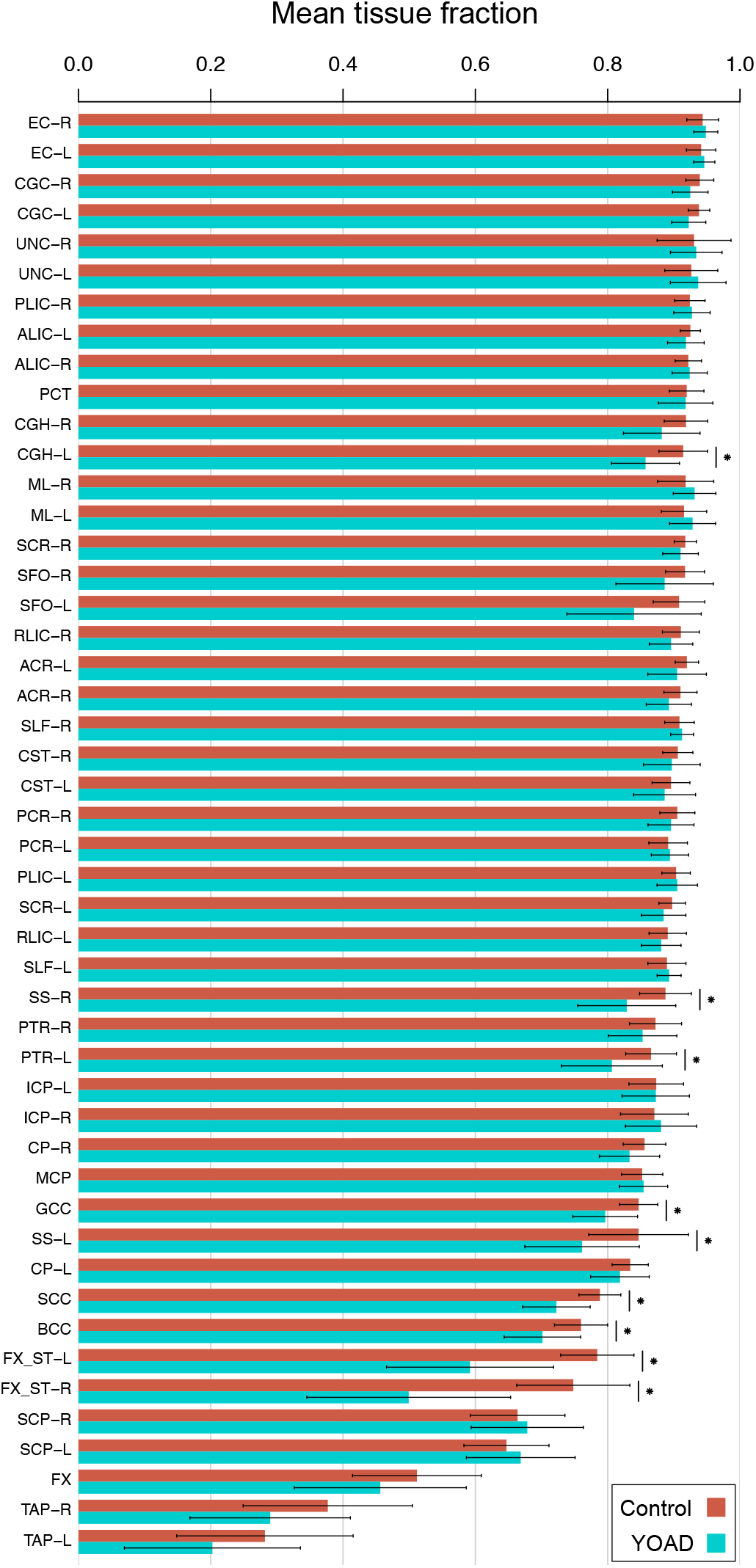
Mean TFs in control and YOAD subjects for each white matter ROI. Bars show the mean ± standard deviation of the mean TF across subjects. ROIs are in decreasing order of mean TF in control subjects from left to right. Bilateral ROIs are ordered adjacently when no significant difference was observed by two-tailed t-test between their mean TFs in control subjects. Horizontal lines with stars denote significantly lower mean TF in YOAD subjects, determined using two-tailed Welch’s t-tests (*p*<0.05 Bonferroni-corrected across ROIs).

### 4.2. Bias in the conventional mean

There is statistically significant evidence of bias in the conventional mean of NODDI metrics for most white matter ROIs (Fig. 4). Those with lower mean TF tend to have higher magnitudes of bias, as expected from equation (6) – the correlation between mean magnitude of bias and the inverse of mean TF is high (*r*>0.94) for both NODDI tissue metrics (NDI, ODI) and cohort group (control, YOAD) combinations [*r*=0.94 for NDI in controls (*p*=6.0×10^-24^), *r*=0.95 for NDI in YOAD (*p*=3.0×10^-25^), *r*=0.99 for ODI in controls (*p*=7.7×10^-41^) and *r*=0.99 for ODI in YOAD (1.1×10^-39^)]. As implied, YOAD subjects’ ROIs with relatively lower mean TF tend to display greater bias.

**Figure 4.**
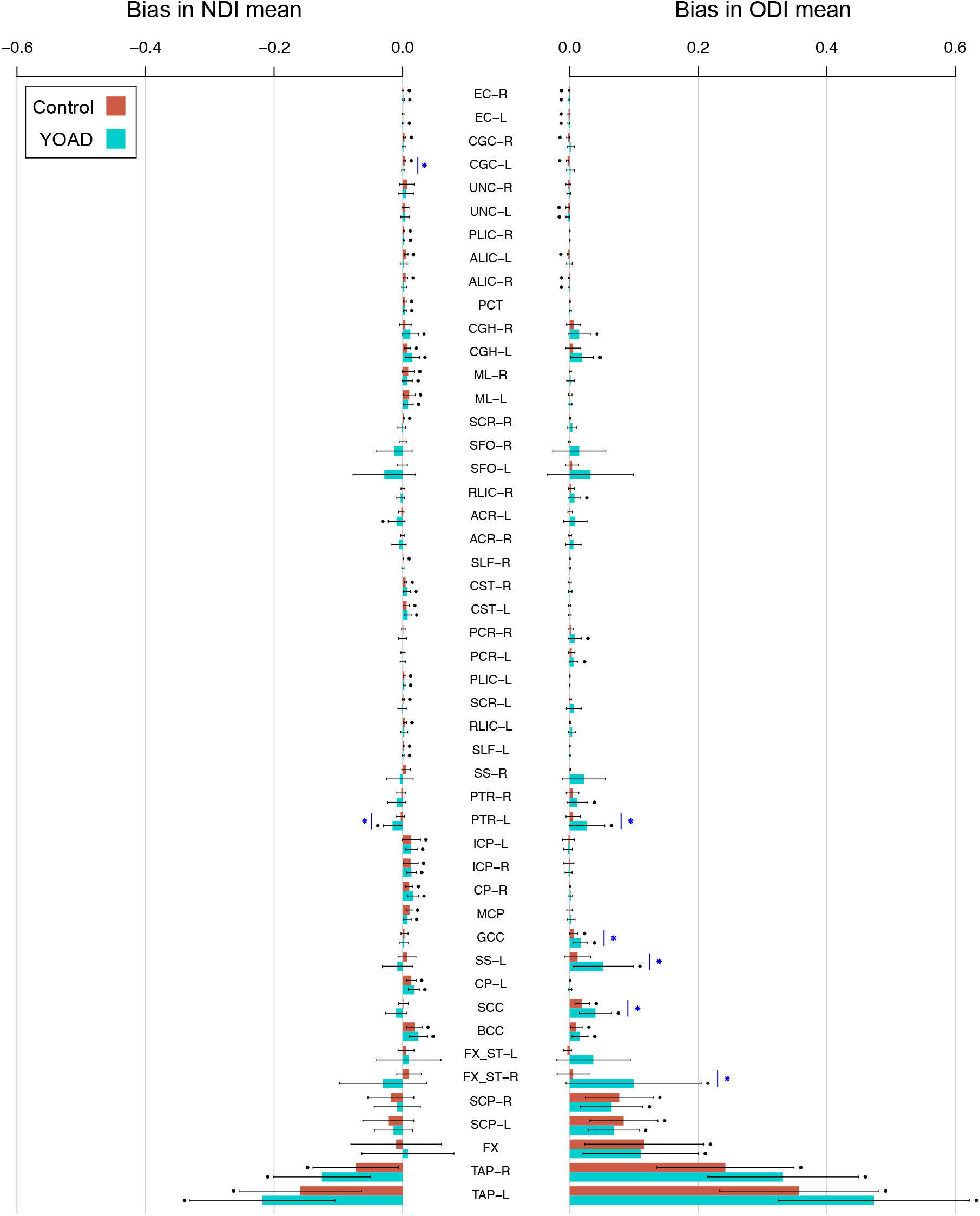
Bias in conventional means for each white matter ROI. Bars show the mean ± standard deviation of bias across subjects. The height of each bar is the average bias across subjects, equal to the bias in the group mean. Black points indicate significant evidence of non-zero bias, determined using two-tailed one sample t-tests (*p*<0.05 Bonferroni-corrected across ROIs). Blue stars indicate significant differences in bias between control and YOAD subjects, determined using two-tailed Welch’s t-tests (*p*<0.05 Bonferroni-corrected across ROIs). ROIs are ordered as in Fig. 3.

**Figure 5.**
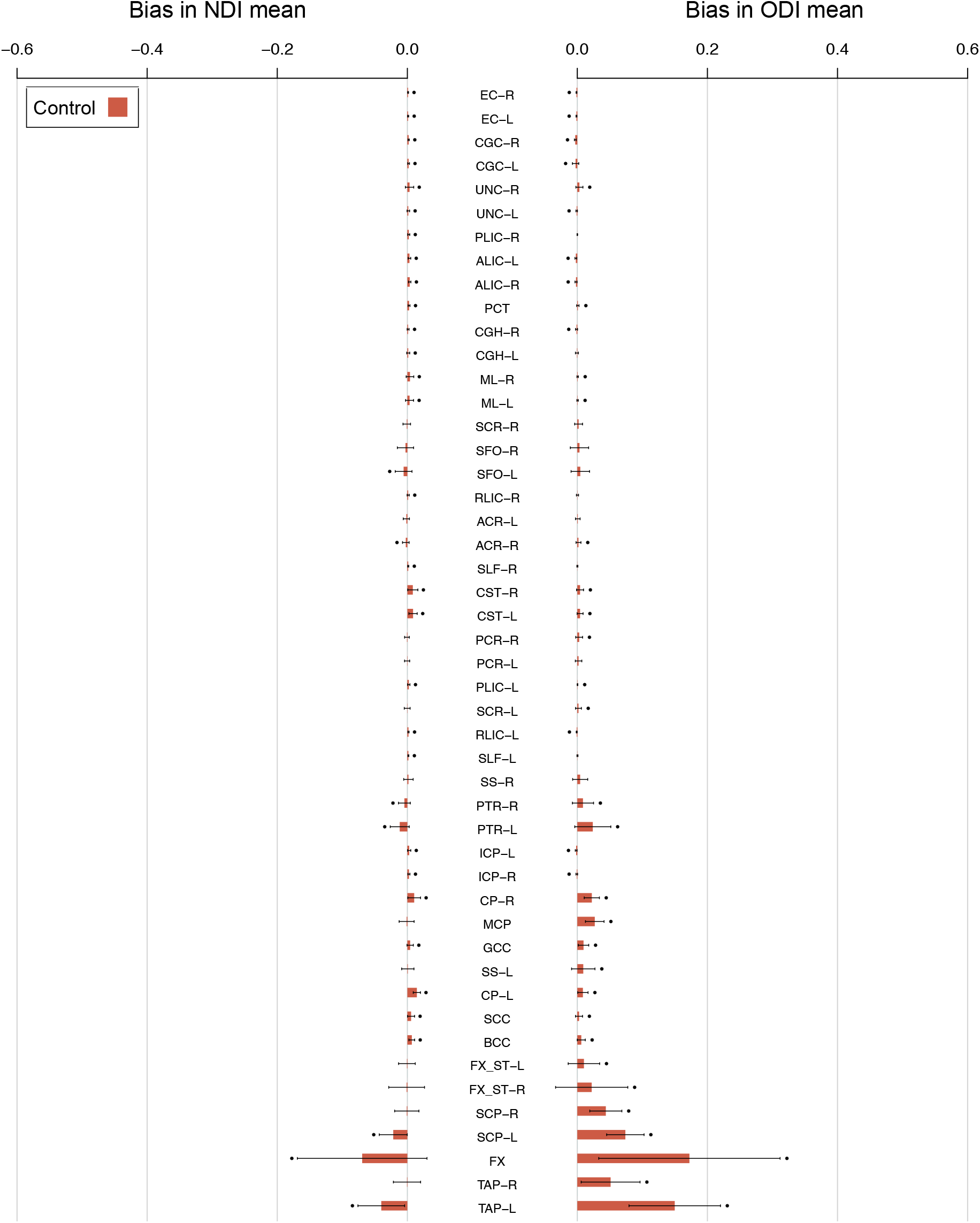
Bias in conventional means for each white matter ROI in higher resolution ADNI data. The height of each bar is the average bias across ADNI control subjects. Black points indicate significant non-zero bias, as in Fig. 4.

Periventricular ROIs tend to have higher magnitudes of bias than non-periventricular ROIs. Significantly higher bias for ODI in control subjects (*p*=0.043) and YOAD subjects (*p*=0.032) is observed for periventricular ROIs, whereas NDI bias tends to be higher in periventricular ROIs for both groups but is not significant (control *p*=0.067, YOAD *p*=0.11).

ODI is positively biased (over-estimated), an effect more pronounced in ROIs with lower mean TF. NDI bias in non-periventricular ROIs tends to be positive (over-estimated), whereas NDI bias in periventricular ROIs shows less trends of directionality. The directionality of bias reflects the sign of the covariance between TF and NDI or ODI within an ROI.

### 4.3. Bias at higher image resolution

As with the relatively lower resolution YOAD data, the correlation between mean magnitude of bias and the inverse of mean TF is high (>0.9) for both NODDI tissue metrics [0.90 for NDI (*p*=8.7×10^-18^) and 0.96 for ODI (3.7×10^-28^)]. When considering the mean across control subjects for each ROI, the magnitude of bias tends to be lower in the higher resolution ADNI data than in the YOAD cohort but is not significant.

### 4.4. Bias in template space ROIs

A similar association between lower mean TF and higher magnitudes of bias, higher bias in periventricular ROIs, and higher bias in the patient group is observed in the template space as with the native space ROIs (Fig. S4, S5). Accordingly, the group mean bias is highly correlated between native and template space ROIs for both groups of subjects (control NDI *r*=0.96, ODI *r*=0.99; YOAD NDI *r*=0.97, ODI *r*=0.99).

A strong positive association (*r*=0.89-0.98, all *p*<5.7×10^-17^ for metric-group combinations) between bias magnitude and inverse of mean TF is observed in template space ROIs, consistent with bias associated with CSF contamination. Periventricular ROIs show significantly greater magnitudes of bias in ODI than non-periventricular ROIs for both cohort groups (*p*<0.05), a trend is also observed for NDI but which is not significant (*p*<0.073). YOAD subjects’ ROIs with their relatively lower mean TF tend to display greater bias. Bias directionality is similar in the template space as in the native space for most ROIs.

Two main differences in patterns of bias are observed in template space: - bias magnitudes tend to be lower for non-periventricular ROIs and bias in NDI conventional means tend to be more consistently negative (under-estimated) in ROIs with lower mean TF.

### 4.5. Group differences comparison

Fig. 6 compares the estimated effect sizes computed using the conventional mean and tissue-weighted mean (Fig. S2, S3). Overall, effect sizes are similar using the two approaches, with only small differences observed for the majority of ROIs. However, those ROIs with large differences in bias between control and YOAD groups (Fig. 4) have large differences in effect sizes. Note that as we are interested in the direct comparison between the tissue-weighted mean and conventional ROI mean, we report unadjusted effect sizes as these are quantitatively simple to interpret. We found that adjusting the NODDI tissue metrics for subtle variations in age and sex between groups had a negligeable influence on the estimated effect sizes (Fig. S6).

**Figure 6.**
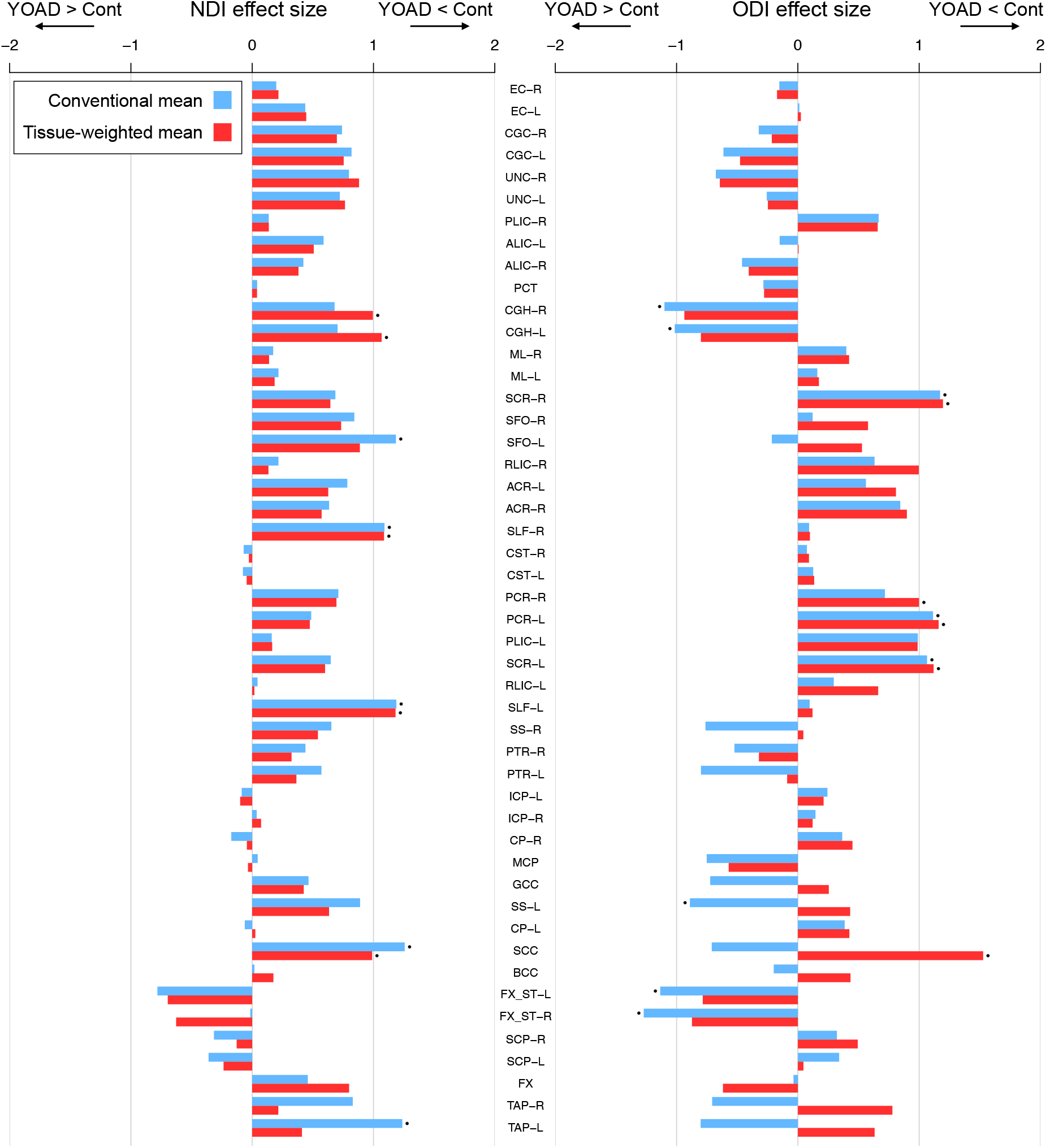
Effect sizes for group differences in NDI (left) and ODI (right) between control and YOAD subjects (Cohen’s d_s_, mean difference contrast of control minus YOAD) using the conventional mean (blue) and tissue-weighted mean (red), for each white matter ROI. Positive effect sizes correspond to lower means in YOAD subjects and negative effect sizes to higher means in YOAD subjects. Points above the bars indicate significant differences between the control and YOAD group as determined by two-tailed Welch’s t-tests on the group means (*p*<0.05 Bonferroni-corrected across ROIs). ROIs are ordered as in Fig. 3.

Using the conventional mean, effect sizes for lower NDI in YOAD compared to controls are over-estimated in comparison to the tissue-weighted mean for most ROIs (28/39 showing lower NDI, Fig. 6), with the largest over-estimation observed for the left and right tapetums (TAP-L and TAP-R) and superior frontal occipital fasciculus (L-SFO) (note that some effect size differences are visually unapparent). In these ROIs, bias is higher in the group means of the YOAD group than in controls. However, effect sizes for lower NDI in YOAD are under-estimated using the conventional mean for 11 ROIs in which the bias is more positive in the YOAD group than in controls, for example in the fornix (FX). This results in a gain of significant group differences for reduced NDI in the left and right hippocampal cingulum (CGH-L and CGH-R) using the tissue-weighted mean.

Effect sizes for lower ODI in YOAD are under-estimated for most ROIs (19/24 with lower ODI) when using the conventional mean (Fig. 6). In some ROIs, such as those of the corpus callosum (GCC, SCC, BCC), effect size directions using the conventional mean are mis-identified as higher ODI in YOAD. These effects are due to higher (more positive) ODI bias in the mean of YOAD groups than in controls. Using the tissue-weighted mean results in a gain of significant group differences for lower ODI in YOAD in the right posterior corona radiata (PCR-R) and splenium of the corpus callosum (SCC) which are absent using the conventional mean. There are also ROIs with over-estimated effect sizes for higher ODI in YOAD using the conventional mean. Significant group differences for higher ODI in YOAD are absent for the left and right hippocampal cingulum, limbs of the fornix (FX_ST-R and FX_ST-L) and left sagittal stratum (SS-L) when using the tissue-weighted mean compared to the conventional mean. Effect size for higher ODI in YOAD increases in the fornix when using the tissue-weighted mean.

## 5. Discussion

This study introduces the tissue-weighted mean, an unbiased method for estimating the mean of NODDI tissue microstructure metrics within an ROI. We observe statistically significant bias in conventional means for most ROIs and an association between higher magnitudes of bias and lower tissue fraction. In addition to the subjects’ native space, bias is observed in images at higher resolution and when warping images to a template. Furthermore, due to its higher magnitude in patients than healthy subjects, the observed bias confounds the estimation of group differences, resulting in effect sizes of conventional means being either over- or under-estimated compared to the tissue-weighted mean.

Bias in conventional means occurs because the contribution of voxels with low TF are over-weighted. Periventricular ROIs, which tend to have lower mean TF consistent with CSF partial volume contamination, have higher magnitudes of bias. These findings suggest that in general the periventricular structures, such as the corpus callosum, are particularly susceptible to bias. Certain small periventricular structures, such as the tapetum and fornix, appear even more susceptible due to the relatively high proportion of their surface bordering CSF and larger surface area to volume ratio. As brain atrophy leads to reduced regional volumes (Vernooij et al 2008, Agosta et al 2011) and enlarged ventricles (Drayer et al 1985), regions that have undergone atrophy may have an even higher propensity for bias, which is supported by findings of higher bias in the YOAD group.

The choice of image analysis space in which to compute ROI means may also influence the observed bias. Previous studies have performed ROI analysis in either the native subject space (e.g., Oishi et al 2009) or template space (e.g., Geng et al 2012). As template space analysis requires transformation and resampling of NODDI maps, it is important to determine the influence that interpolation and distortion associated with image transformation has on the patterns of bias. When warping NODDI metrics to a template, overall patterns of bias are similar. This indicates that analysis of regional NODDI tissue metrics in either space should use the tissue-weighted mean to reduce estimation bias. However, marginally lower magnitudes of bias are observed in some ROIs. Although the differences are small, this suggests that the computational processing of images prior to ROI analysis (i.e., warping to a population template) can have an impact on the magnitude of bias, and that previous studies of ROIs in template space may have experienced lower magnitudes of bias than those in native space.

Bias persisted in conventional ROI means derived from DWI at more standard image resolutions (2mm isotropic vs. 2.5mm isotropic voxels in YOAD). In the higher resolution ADNI 3 data, despite ROIs containing twice the number of voxels (125 per 1000mm^3^ in 2mm isotropic data vs. 64 per 1000mm^3^ in 2.5mm isotropic data) and therefore having a substantially lower proportion of voxels with CSF partial volume, the overall magnitude of bias is not significantly lower. This demonstrates that bias observed in the YOAD cohort is not purely a result of the DWI data having relatively lower resolution, and that bias can affect images at resolutions which are now standard in neuroimaging research (Scahill et al 2020). Ongoing large-scale population studies such as ADNI (Weiner et al 2017) and UK Biobank (Miller et al 2016, Alfaro-Almagro et al 2018), which make open access datasets of biomarkers that include summary metrics of microstructure in white matter ROIs, may benefit from more accurate estimates by applying the tissue-weighted mean.

We expect that bias can affect studies in mice as well as in humans, based on their similar relative resolution and ROI positions relative to the ventricles. While it is true that image resolution for mice is considerably higher than for humans, it is the voxel size relative to the size of anatomical structures that is important when comparing across species. Indeed, this relative image resolution is similar in mouse and human studies. Given the reported brain volumes of mice (Badea et al 2007) and humans (Hofmann 2014, Im et al 2008) are ~509 and ~1,400,000 mm^3^ respectively, both have on the order of tens of thousands of voxels per brain volume at (200 x 200 x 500) μm^3^ (Colgan et al 2016) and (2.5 x 2.5 x 2.5) mm^3^ resolutions. In fact, the human brain has ~90,000 voxels per brain volume compared to the mouse which has ~25,000. The anatomical location of ROIs with respect to CSF is also similar among the white matter structures. For example, both human and mice have the midsagittal portion of the corpus callosum bordering the lateral ventricles at the aforementioned imaging resolutions.

Bias in conventional means impacts estimation of group differences, with effect sizes computed using the conventional mean being either under- or over-estimated (Fig. 6). The magnitude of effect size mis-estimation is region-dependent, with the ROIs having higher differences in bias magnitudes between groups also having higher differences in effect sizes. Effect sizes in structures containing substantial partial volume tend to be more severely affected. These results suggest that bias in conventional means can confound estimation of effect sizes and alter findings of significant group differences. Inference of group differences in regional microstructure using conventional ROI means of NODDI tissue metrics may be influenced by bias, particularly those in periventricular regions and those undergoing atrophy. Firstly, some true disease effects can be masked by bias, as evidenced by ROIs that have higher effect sizes when using the tissue-weighted mean compared to the conventional mean. Secondly, the conventional mean may over-estimate the effect sizes for some ROIs.

Previous work has addressed the problem of CSF partial volume effects, where the voxel-wise metrics are confounded by CSF partial volume (Vos et al 2011, Metzler-Baddeley et al 2012) by analysing a subset of voxels with minimal partial volume (Zhang, S. & Arfanakis, Liu et al 2011), applying streamline density-based weighting when computing the mean (Lynch et al 2020), or using the median across the ROI instead of the conventional mean (Lewis et al 2018). We emphasise that the tissue-weighted mean aims to address a different problem – that of bias in conventional ROI means that arises after voxel-wise partial volume effects have been accounted for. Nevertheless, as the median might provide a viable alternative method to compute the mean metrics across an ROI, as it is typically not affected by the outlier values that partial volume may cause, we compared the performance of the median to the conventional mean in terms of its ability to reduce bias with respect to the tissue-weighted mean. We found that bias in the median is of a similar magnitude to that of the conventional mean, suggesting that the median is not a substitute for the tissue-weighted mean (Fig. S7).

The tissue-weighted mean is likely applicable to a wide class of DWI models, anatomical locations, research hypothesis and study groups. The method can be applied to tissue metrics derived from any multi-compartment models that estimates the CSF volume fraction, such as the free water elimination (FWE) method (Pasternak et al 2009). The regions that experience CSF contamination are not restricted to periventricular locations, but include other structures that border CSF, such as the neocortical grey matter which is adjacent to the sub-arachnoid space. Aside from neurodegeneration, the tissue-weighted mean can be applied to other diseases which feature ventricular enlargement, such as hydrocephalus, including normal appearing hydrocephalus (Vanneste et al 2000, Corkill et al 2003). Beyond CSF contamination, the tissue-weighted mean can be used under an alternative hypothesis where the free water fraction (CSF volume fraction) corresponds to interstitial free water, such as is the case in inflammation induced vasogenic oedema (Palacios et al 2020). Furthermore, the concept of the tissue-weighted mean is naturally extendable to other descriptive statistics of ROIs that summarise tissue microstructure information across voxels, such as the variance, covariance and regression coefficients.

In this exemplar application of the tissue-weighted mean, the inclusion of YOAD allows us to make tentative assessment of the regional microstructure correlates of the disease. We observe a concordance between significant group differences and expected regional pathology, demonstrating that the tissue-weighted mean is sensitive to biologically plausible disease effects. For instance, the tissue-weighted mean had significantly lower NDI, consistent with tissue neurodegeneration, in the left and right hippocampal cingulum (CHG-L and CGH-R) (Fig. 6, S2), which were absent using the conventional mean. These regions form hippocampal connections involved in memory processing (Nakata et al 2009), a brain function associated with symptoms of Alzheimer’s disease and YOAD (Rossor et al 2010).

## 6. Conclusion

This study shows bias in conventional ROI means is highly prevalent and particularly affects periventricular regions where partial volume due to CSF contamination is higher. This bias confounds group difference metrics, suggesting inferences in cohorts with brain atrophy or different ventricle sizes can be influenced by bias. The proposed tissue-weighted mean provides unbiased estimation of regional mean tissue metrics and can be derived from DWI models that estimate CSF contamination and tissue microstructure. It can be applied to accurately identify disease effects in future studies of white matter neurodegeneration, especially for periventricular regions and for other brain tissues prone to CSF contamination, such as cortical grey matter. This may provide additional insight into associations between brain microstructure and aging, development and neurodegeneration.

## Acknowledgments

CP and GZ were funded by the Wellcome Trust (Collaborative Award 200181/Z/15/Z). TV was funded by an Alzheimer’s Research UK PhD scholarship (ARUK-PhD2018-009). MB was supported by a Fellowship award from the Alzheimer’s Society, UK (AS-JF-19a-004-517). MB’s work was also supported by the UK Dementia Research Institute which receives its funding from DRI Ltd, funded by the UK Medical Research Council, Alzheimer’s Society and Alzheimer’s Research UK. IM was supported by Alzheimer’s Research UK (ARUK-PG2014-1946, ARUK-PG2017-1946) and the Wolfson Foundation (PR/ylr/18575). DLT was supported by the UCL Leonard Wolfson Experimental Neurology Centre (PR/ylr/18575), UCLH NIHR Biomedical Research Centre and the Wellcome Trust (Centre award 539208). JMS acknowledges the support of the National Institute for Health Research University College London Hospitals Biomedical Research Centre, Wolfson Foundation, Alzheimer’s Research UK, Brain Research UK, Weston Brain Institute, Medical Research Council, British Heart Foundation, UK Dementia Research Institute and Alzheimer’s Association. DMC was supported by the UK Dementia Research Institute which receives its funding from DRI Ltd, funded by the UK Medical Research Council, Alzheimer’s Society and Alzheimer’s Research UK, as well as Alzheimer’s Research UK (ARUK-PG2017-1946) and the UCL/UCLH NIHR Biomedical Research Centre. We would also like to acknowledge Prof. Nick Fox who is a senior NIHR investigator for his role in conceiving the initial YOAD study preceding this work.

The authors would like to thank all research participants who made this study possible, as well as Alzheimer’s Research UK and Iceland Foods Charitable Foundation for funding the Young-Onset Alzheimer’s disease study. The Dementia Research Centre is supported by Alzheimer’s Research UK, Brain Research Trust, and The Wolfson Foundation. They also thank Kirsty Lu, Amelia Carton, Timothy Shakespeare, Keir Yong, Aida Suarez Gonzalez and Silvia Primativo for assistance with neuropsychology assessments.

## Appendix

**Table A1.** JHU atlas white matter ROI location, abbreviations and region names.

| Periventricular |  |
| --- | --- |
| MCP | Middle cerebellar peduncle |
| GCC | Genu of corpus callosum |
| BCC | Body of corpus callosum |
| SCC | Splenium of corpus callosum |
| FX | Fornix |
| ML-R | Medial lemniscus right |
| ML-L | Medial lemniscus left |
| SCP-R | Superior cerebellar peduncle right |
| SCP-L | Superior cerebellar peduncle left |
| CP-R | Cerebral peduncle right |
| CP-L | Cerebral peduncle left |
| ALIC-R | Anterior limb of internal capsule right |
| ALIC-L | Anterior limb of internal capsule left |
| RLIC-R | Retrolenticular part of internal capsule right |
| RLIC-L | Retrolenticular part of internal capsule left |
| ACR-R | Anterior corona radiata right |
| ACR-L | Anterior corona radiata left |
| PCR-R | Posterior corona radiata right |
| PCR-L | Posterior corona radiata left |
| SS-R | Sagittal stratum right |
| SS-L | Sagittal stratum left |
| EC-R | External capsule right |
| EC-L | External capsule left |
| CGH-R | Cingulum (hippocampus) right |
| CGH-L | Cingulum (hippocampus) left |
| FX-ST-R | Fornix or stria-terminalis right |
| FX-ST-L | Fornix or stria-terminalis left |
| TAP-R | Tapetum right |
| TAP-L | Tapetum left |
| Non-Periventricular |  |
| PCT | Pontine crossing tract |
| CST-R | Corticospinal tract right |
| CST-L | Cortical spinal tract left |
| ICP-R | Inferior cerebellar peduncle right |
| ICP-L | Inferior cerebellar peduncle left |
| PLIC-R | Posterior limb internal capsule right |
| PLIC-L | Posterior limb internal capsule left |
| SCR-R | Superior corona radiata right |
| SCR-L | Superior corona radiata left |
| PTR-R | Posterior thalamic radiation right |
| PTR-L | Posterior thalamic radiation left |
| CGC-R | Cingulum right |
| CGC-L | Cingulum left |
| SLF-R | Superior longitudinal fasciculus right |
| SLF-L | Superior longitudinal fasciculus left |
| SFO-R | Superior frontal occipital fasciculus right |
| SFO-L | Superior frontal occipital fasciculus left |
| UNC-R | Uncinate fasciculus right |
| UNC-L | Uncinate fasciculus left |

