## Supplementary Material for "Not all voxels are created equal: reducing estimation bias in regional NODDI metrics using tissue-weighted means"

*S.1. White matter regions of interest*

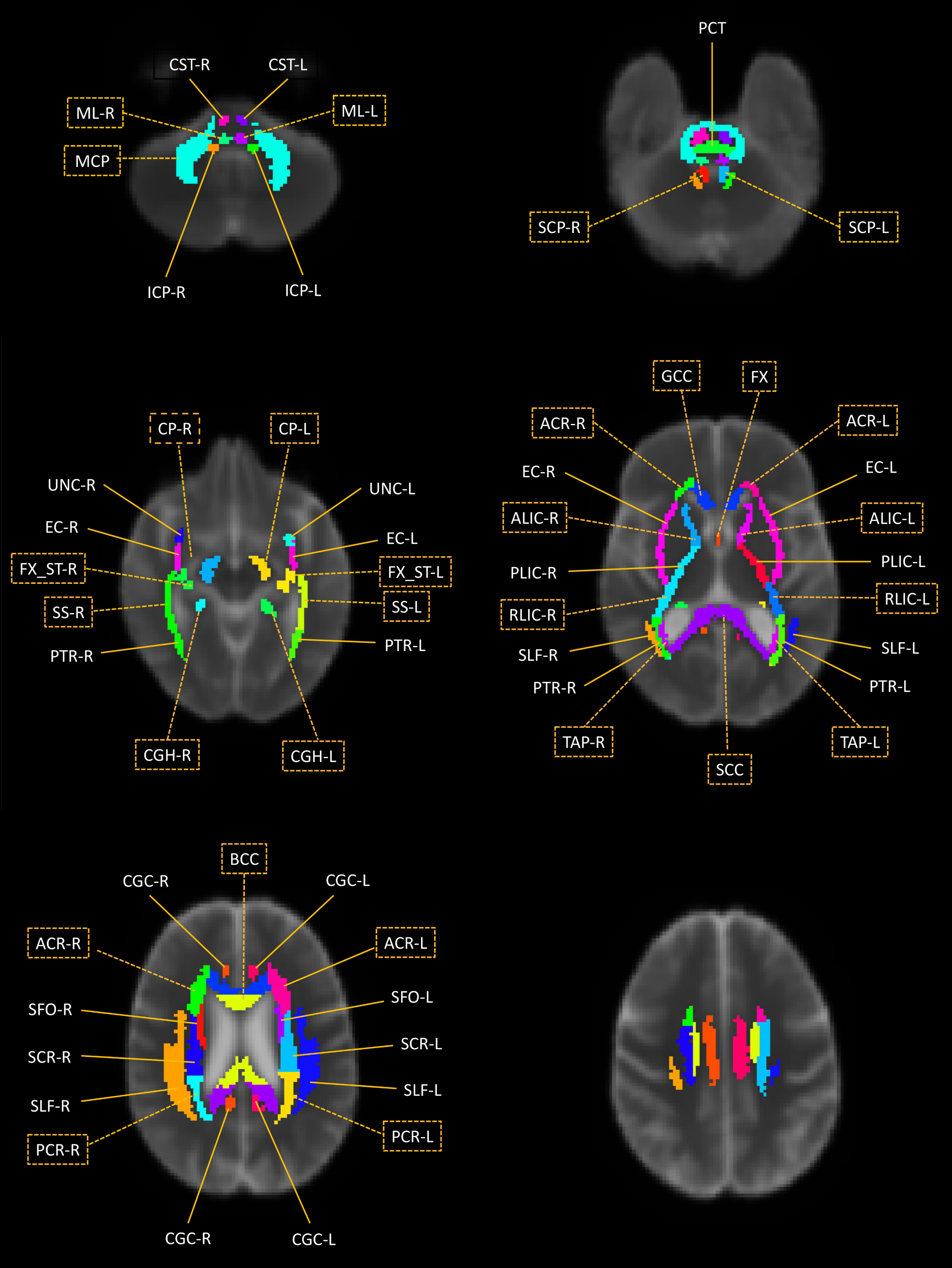

**Figure S1.** White matter ROIs overlaid on the mean diffusivity map of the population template. ROI abbreviations are listed in Table A1. Yellow boxes surround abbreviations of periventricular ROIs.

*S.2. YOAD subject characteristics*

The YOAD study (Slattery et al 2017) recruited 24 controls and 45 YOAD patients matched for age and sex from a specialist cognitive disorders clinic between 2013 and 2015. Diagnosis of YOAD was based on published consensus criteria for probably AD (McKhann et al 2011) and symptom onset < 65 years of age. Out of the original 69 recruited subjects we analysed 21 control subjects and 30 YOAD patients that passed image quality control. 18 were excluded due to motion artefacts or processing failures. Specifically, 1 was excluded due to excessive T1w motion, 3 due to image processing failures of T1w images, 7 due to motion and processing failures in DTI sequences and 7 due to motion and processing failures in NODDI sequences, as described (Veale et al 2021). Study participants demographic characteristics are shown in Table S1.

|  | Control n=20 | | YOAD n=31 | | *p* |
| --- | --- | --- | --- | --- | --- |
|  | Mean | SD | Mean | SD |  |
| Age | 60.5 | 5.7 | 61.4 | 4.6 | 0.5^a^ |
| Sex M:F | 9:12 | - | 13:17 | - | 0.2^b^ |
| Education | 16.7 | 3.2 | 15.1 | 2.8 | 0.1^c^ |
|  |  |  | N |  |  |
| APOE +ve | - | - | 16 | - | - |

**Table S1.** Study participants’ demographic characteristics. Key: SD, standard deviation; M, male; F, female; APOE, apolipoprotein E.

^a^ Two-tailed T-test

^b^ Two-tailed Fischer exact test

^c^ Mann-Whitney U test

*S.3. ADNI subject characteristics*

Imaging data for the 75 healthy control subjects were obtained from ADNI 3 in October 2020. The subjects have a mean age of 72.9 years old and a standard deviation of 7 years. 26 out of the 75 were male.

*S.4. Conventional and tissue-weighted means*

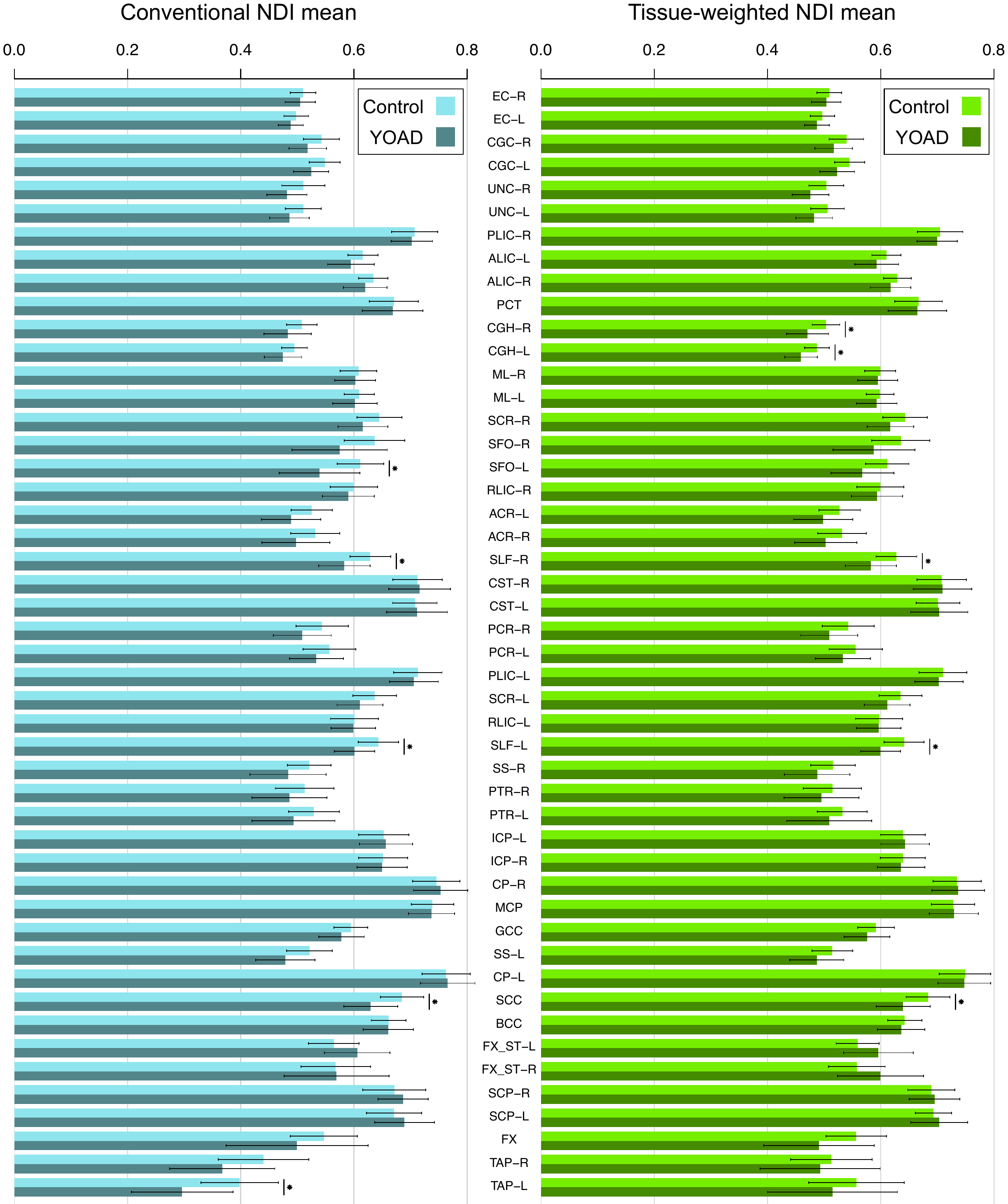

**Figure S2.** Conventional mean and tissue-weighted mean NDI in control and YOAD subjects. Bars show the group mean ± standard deviation. Stars show a significant group differences between control and YOAD ROI means (*p*<0.05 Bonferroni corrected), as determined using two-tailed Welch’s t-tests. Regions are ordered as in Figure 3.

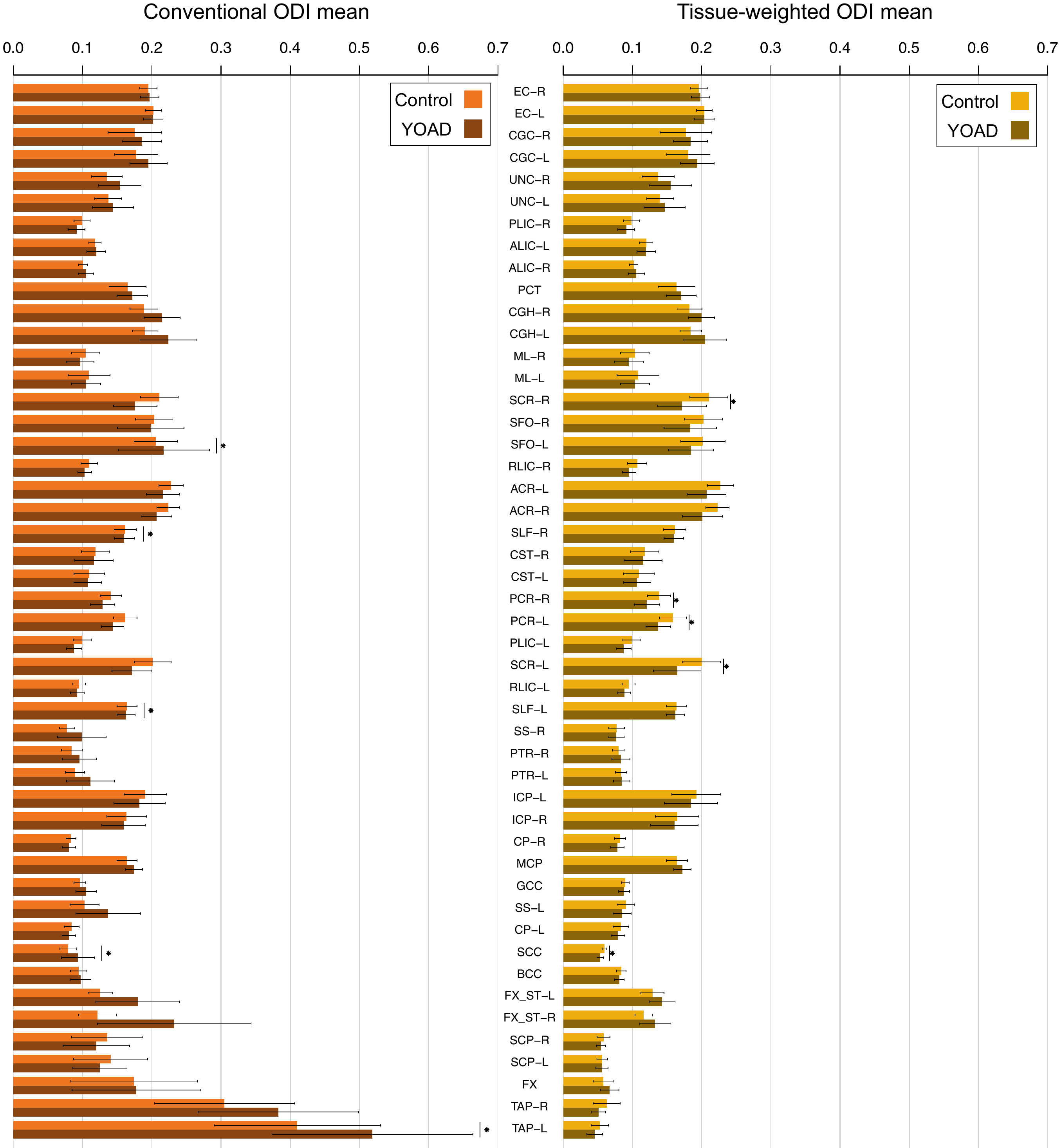

**Figure S3.** Conventional mean and tissue-weighted mean ODI in control and YOAD subjects. Figure interpretation follows that of Fig. S2.

*S.5. Bias in template space ROIs*

*S.5.1. Mean tissue fractions*

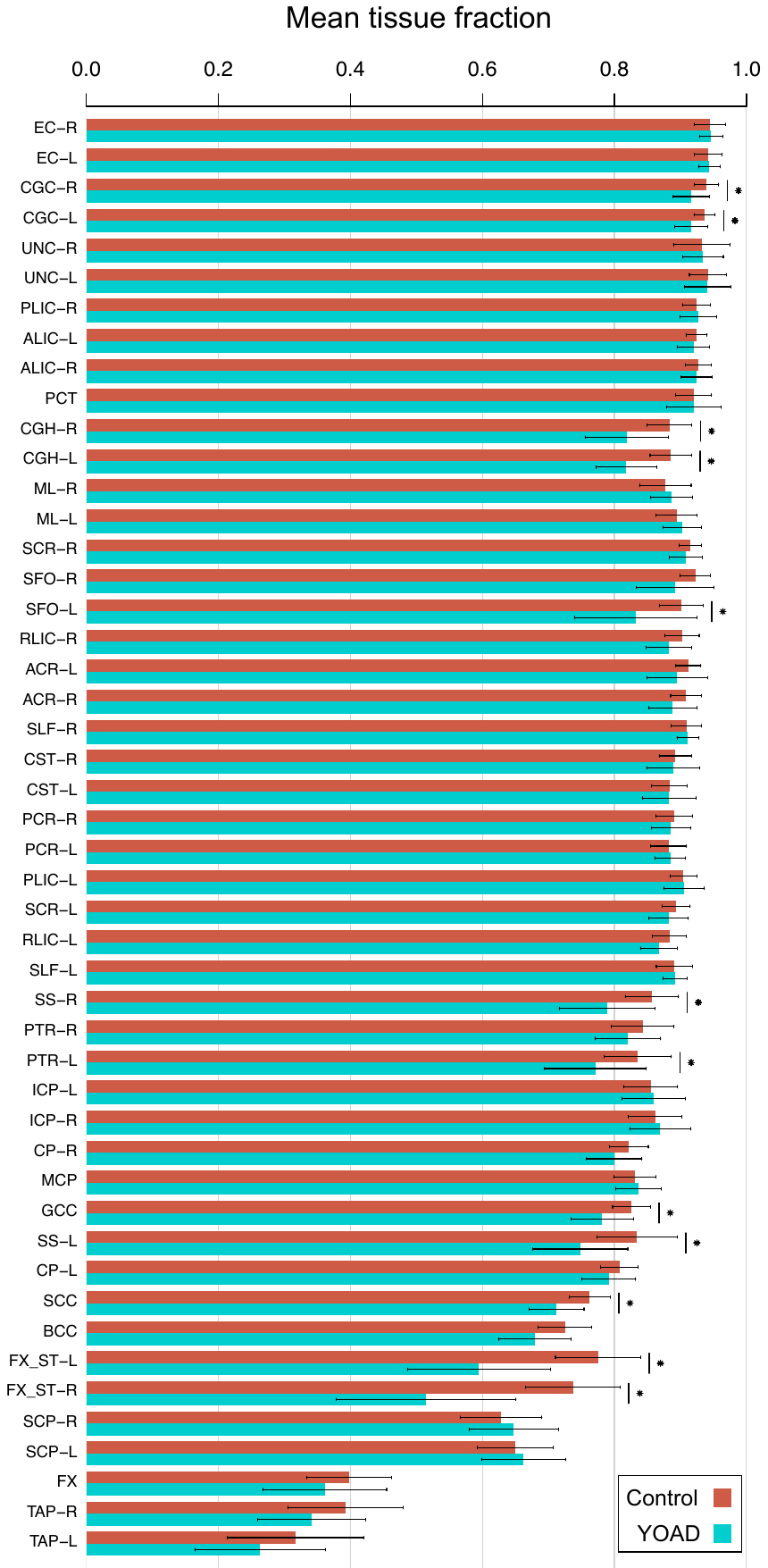

**Figure S4.** Mean TFs in control and YOAD subjects for each white matter ROI in template space. Figure interpretation is the same as in Fig. 3.

*S.5.2. Bias in conventional means*

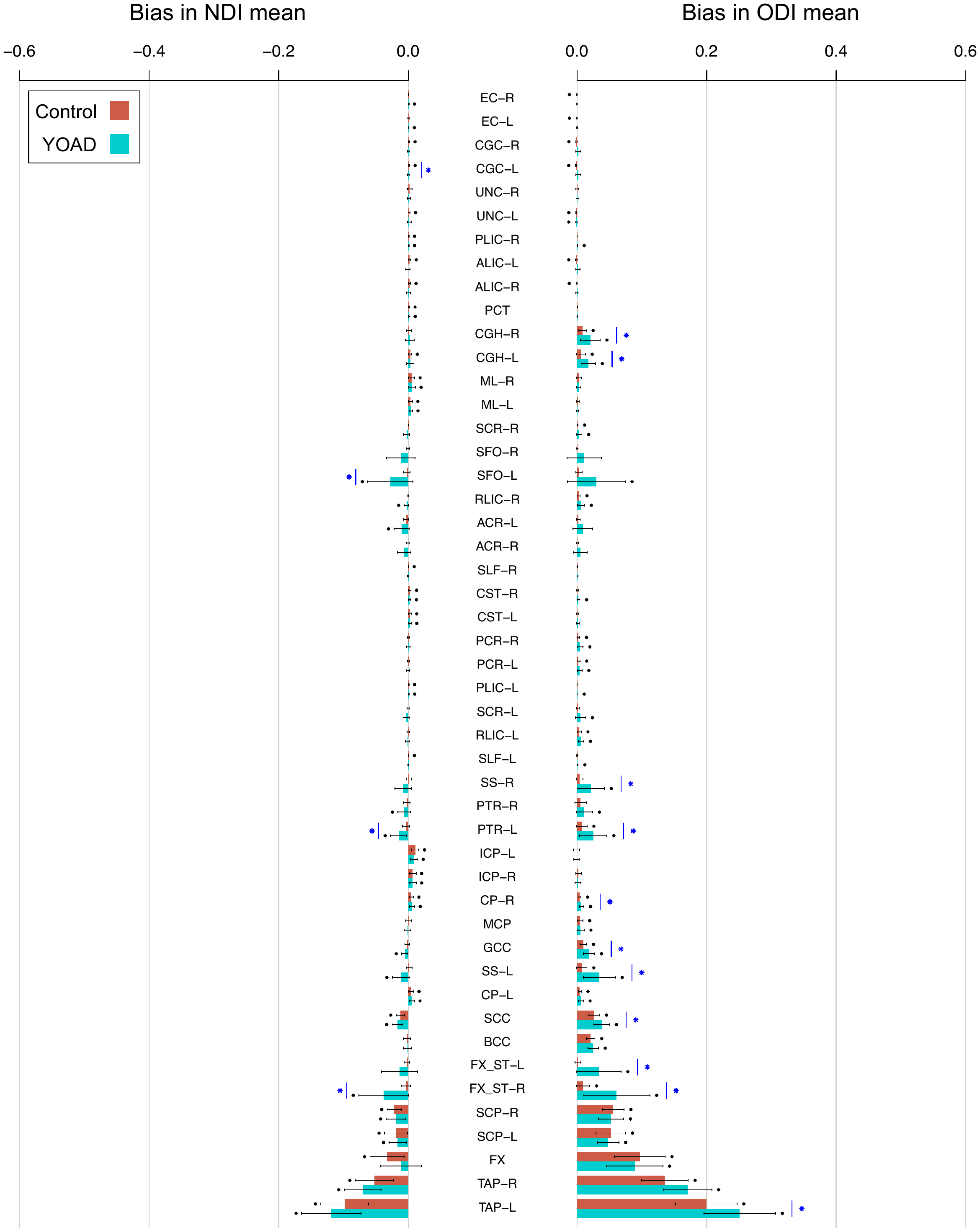

**Figure S5.** Bias in conventional means for each white matter ROI in template space. Figure interpretation is as in Fig. 4.

*S.6. Group differences comparison adjusting for covariates*

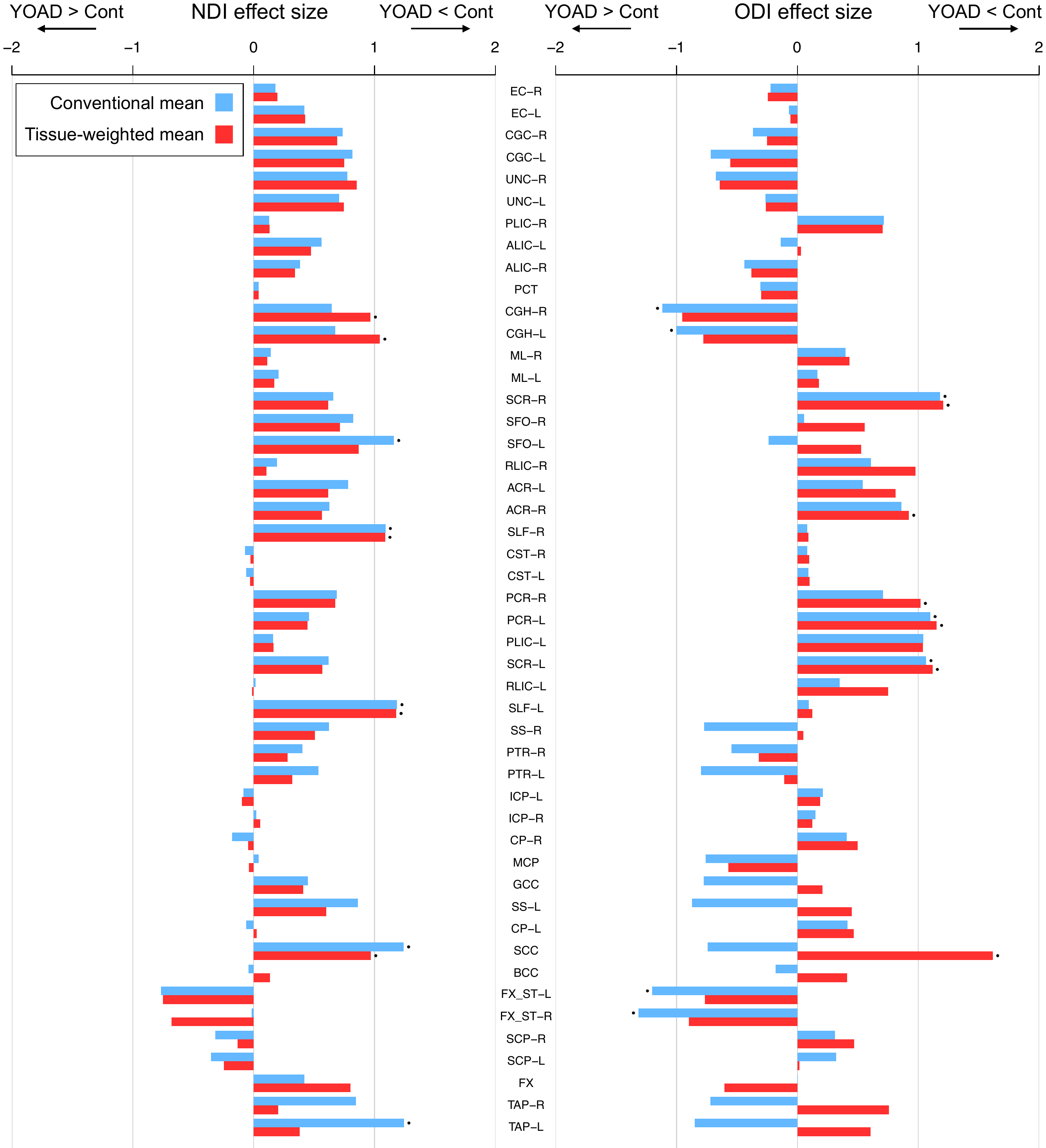

**Figure S6.** Effect sizes for group differences in covariate-adjusted NDI (left) and ODI (right) using the conventional mean and tissue-weighted mean. NDI and ODI were adjusted for age and sex using a linear model. Points above the bars show a significant difference between control and YOAD ROI means. ROI analysis was performed in the native subject space. ROIs are ordered as in Fig. 3.

*S.7. Bias in ROI median*

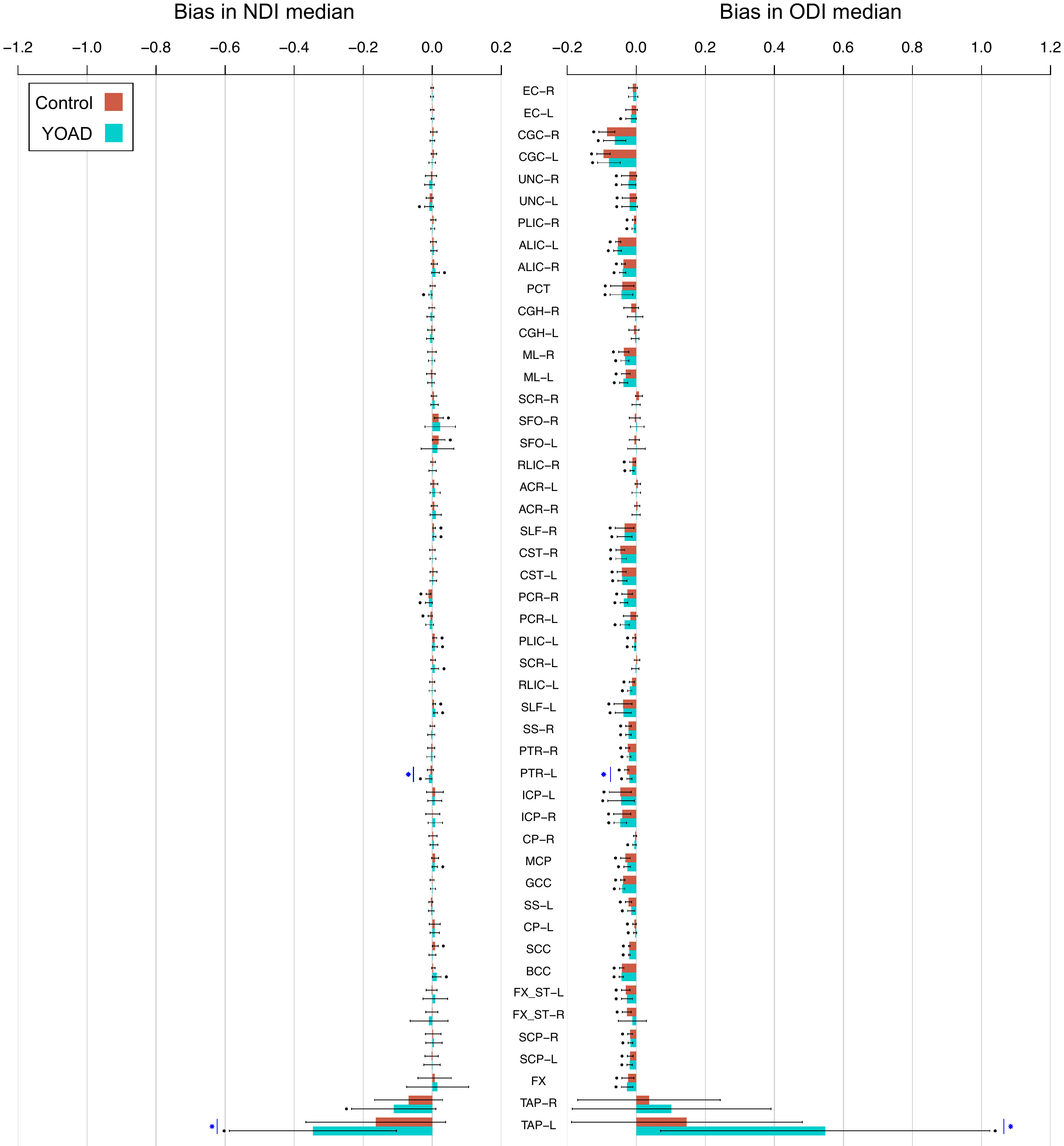

**Figure S7.** Bias in the median for each white matter ROI in native subject space. Figure interpretation is as in Fig. 4.
